# The Embodied Nature of Well-Timed Behavior

**DOI:** 10.1101/716274

**Authors:** Mostafa Safaie, Maria-Teresa Jurado-Parras, Stefania Sarno, Jordane Louis, Corane Karoutchi, Ludovic F. Petit, Matthieu O. Pasquet, Christophe Eloy, David Robbe

**Author notes:** Co-first authors (Mostafa Safaie, Maria-Teresa Jurado-Parras).

## Abstract

How animals adapt their movements to take advantage of behaviorally-relevant time intervals is not well understood, especially in the supra-second timescale. It has been proposed that motor timing depends on the emergence of self-sustained dynamics across ensembles of neurons. Alternatively, evidence from operant conditioning suggests that animals can develop motor routines to adapt their behavior to fixed temporal constraints. But it is unclear whether animals can accurately time their behavior without the help of motor routines. To address this issue, we used a task in which rats, freely moving on a motorized treadmill, could obtain a reward if they approached it after a fixed interval. Most animals took advantage of the treadmill length and its moving direction to develop, by trial-and-error, a unique motor routine whose execution resulted in the precise timing of their reward approaches. Noticeably, when proficient animals occasionally failed to follow this routine, the timing of their reward approaches was systematically poor. In a second step, we trained naive animals in modified versions of the task specifically designed to prevent the development of this motor strategy. Compared to rats trained in the first protocol, these animals never reached a comparable level of timing accuracy. We conclude that motor timing critically depends on the ability of animals to develop motor routines adapted to the structure of their environment. Our work also suggests that self-sustained neuronal activity alone may not be sufficient to support motor timing, at least in the supra-second timescale.

## Introduction

The ability of animals to adapt their behavior to periodic events is critical for survival, as the appearance of a sensory cue can predict the timing of food availability, predator attack or mating opportunity [1–4]. Understanding the mechanisms underlying this ability is challenging because unlike sensory modalities (vision, olfaction, audition, …), time is not a material entity and animals are not equipped with a sense organ for time perception. It has been postulated that animals use a dedicated internal clock to estimate the duration of behaviorally-relevant time intervals and sensory cues and produce well-timed movements or take adaptive decisions [5–10]. However, time is a critical parameter for a wide range of behavioral functions (sensory detection, memory, motor control, attention, response to threats) that engage distinct brain regions. In addition, temporal representations are intrinsic to the activity of ensembles of neurons (i.e., neuronal population activity dynamically evolves in time [11]). Thus, the ability of animals to judge the duration of sensory stimuli or to produce movements according to temporal constraints may emerge from the dynamics of task neuronal populations, rather than a time-dedicated internal clock [11, 12]. More specifically, it has been proposed that the well-timed production of movements (i.e., motor timing) could depend on the self-sustained population dynamics that naturally emerges from the activity of recurrently connected neurons [13]. This time-varying emergent signal would be triggered by a cue signalling the beginning of the interval to time and, at the end, will be read by output (motor) neurons controlling movement generation. Still, whether self-sustained neuronal activity *alone* can reliably produce well-timed motor responses independently of external signals (e.g., sensory stimuli appearing during the interval or sensorimotor feedback triggered by movements) is not resolved [13], an issue that is especially relevant for supra-second long intervals or delays.

Interestingly, early investigations using a variety of supra-second long motor timing tasks reported that animals often develop stereotyped chains of actions between the operant responses delivering the reward [14–17]. These so-called collateral behaviors (sometimes referred to as superstitious, adjunctive or interim behaviors) have been observed in a wide range of species, including humans, but, as their names indicate, it is believed that they do not contribute to motor timing per se [17]. Indeed, the duration and order of the collateral behaviors can substantially vary during supra-second long intervals, making them relatively unreliable *external* clocks. Consequently, in the few timing theories that considered collateral behaviors, their duration and transition times were assumed to be largely determined by some sort of internal clocks [18, 19] or habitual processes [20, 21].

Recently, the continuous video monitoring of rats performing different timing tasks demonstrated that proficient animals developed highly stereotyped motor routines [22–24], raising again the possibility that timing could be facilitated by the execution of routines or motor sequences whose durations match the task temporal constraints. Still, it is possible that animals specifically rely on self-sustained neuronal activity when experimental conditions prevent the usage of stereotyped motor strategies. In other words, it is unclear whether animals can accurately time their behavior without the help of motor routines. Moreover, even if one assumes that motor routines do facilitate motor timing, it could still be argued that they are themselves primarily driven by self-sustained neuronal activity.

To address these questions, we challenged rats in a task taking place on a 90 cm-long motorized treadmill, in which animals had to wait for a 7 s long time interval (or delay) before approaching a reward port located at the front of the treadmill. We observed that rats took advantage of the task parameters (treadmill speed, direction and length) to learn, by trial and error, a simple motor routine whose execution resulted in their front-back-front trajectory on the treadmill and an accurate timing of their reward approaches. By manipulating the duration of the waiting time, the speed of the treadmill (its magnitude and reliability across trials), or interfering with the initiation of this simple motor routine, we created conditions that prevented its development or usage. In these conditions, rats were always less accurate and thus seemed unable to rely on a purely internal mechanism. Critically, we observed that accurate timing emerged only when animals could take advantage of salient external features of the environment that served as a scaffold for the execution of the motor routine. We conclude that the production of well-timed behaviors by rodents critically depends on their ability to develop motor routines adapted to the structure of their environment. Our work also suggests that in supra-second motor timing tasks in which animals cannot rely on motor routines, self-sustained neuronal activity alone may not be sufficient to produce well-timed behaviors.

## Results

To investigate how animals adapt their behavior to temporal regularities in their environment, we challenged Long-Evans rats in a treadmill-based behavioral assay that required them to wait for 7 s before approaching a “reward area”. The treadmill was motorized, surrounded by 4 walls and its length (distance between the front and back walls) was 90 cm. The front wall was equipped with a device delivering rewards (a drop of sucrose solution) and an infrared beam, located at 10 cm of this device, defined the limit of the reward area. (Figure 1a). Animals were first familiarized with the apparatus and trained to lick drops of the sucrose solution delivered every minute while the treadmill was immobile (see Methods). Then, rats were trained once a day (Mondays to Fridays) for 55 minutes in the proper waiting task. Each daily session contained ∼130 trials interleaved with resting periods of 15 s (intertrial, motor off). Each trial started by turning the treadmill motor on at a fixed speed of 10 cm/s. The conveyor belt moved toward the rear of the treadmill (Figure 1a). The animals’ entrance time in the reward area (*ET*, detected by the first interruption of the infrared beam in each trial) relative to a goal time (GT, 7 s after motor onset) defined 3 types of trials. Trials in which animals entered the reward area after the GT were classified as correct (7 ≤ *ET* < 15, Figure 1b). Trials in which animals entered the reward area before the GT were classified as error (1.5 ≤ *ET* < 7, Figure 1c). If in 15 s an animal had not interrupted the infrared beam, the trial ended and was classified as omission (Figure 1d). Interruptions that occurred during the first 1.5 s (*ET* < 1.5) were ignored to give the opportunity to the animals to leave (passively or actively) the reward area at the beginning of each trial. Additionally, the exact value of the *ET* determined a reward/punishment ratio. The volume of the sucrose solution delivered, increased linearly for *ET* values between 1.5 s (minimal reward) and GT (maximal reward) and decreased again between GT and 15 s (end of trial). A penalty period of extra running started when the animals erroneously crossed the infrared detector before GT (1.5 ≤ *ET* < 7) and its duration varied between 10 s and 1 s, according to the error magnitude (Figure 1c, inset). In addition, to progressively encourage the animals to enter the reward area after the GT, the smallest *ET* value that triggered reward delivery was raised across sessions, according to each animal’s performance, until it reached the GT (Figure 1b, inset and see Methods for details). Thus, to maximize reward collection and minimize running time, animals should cross the infrared beam just after the GT.

**Figure 1:**
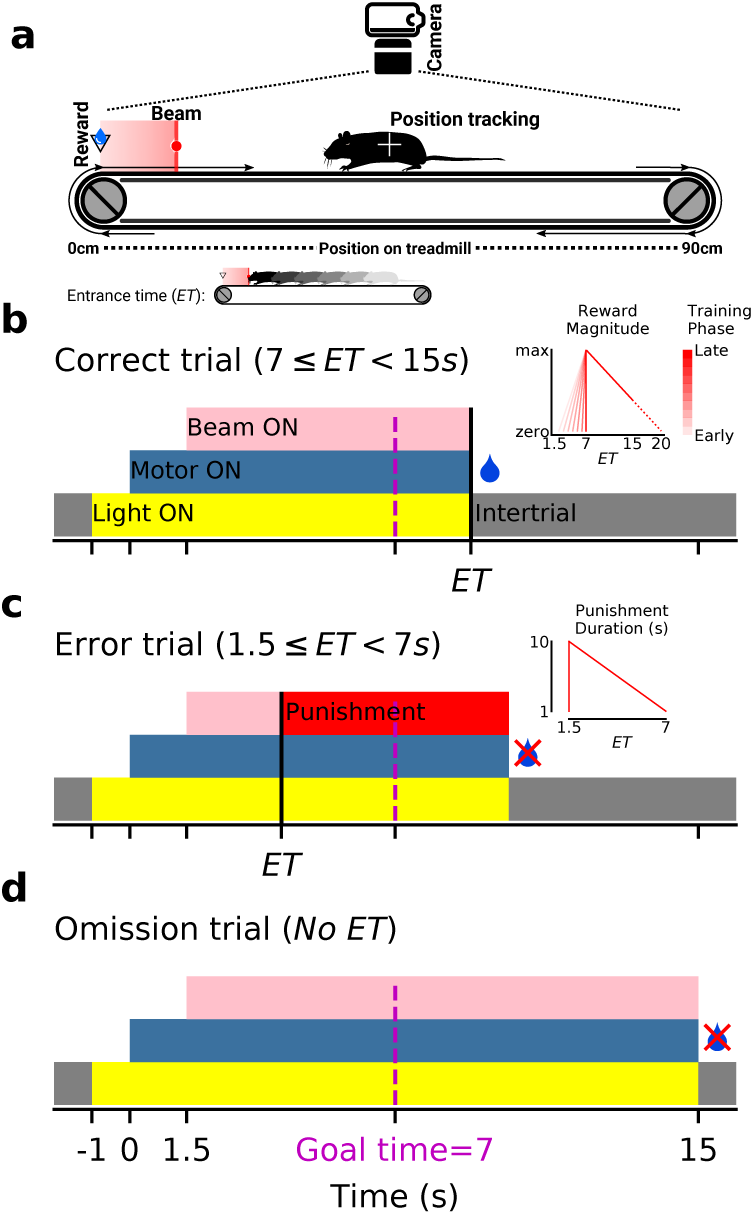
Treadmill task and trial types. **a)** Rats were enclosed on a motorized treadmill. The infrared beam placed at 10 cm of the reward port marked the beginning of the reward area (pink shaded area). During each trial, the belt pushed the animals away from the reward area and the first infrared beam interruption defined the reward area entrance time (*ET*). During trials and intertrials, the animals’ position was tracked via a ceiling-mounted video camera. **b)** Schematic description of a rewarded correct trial. *Inset*: the magnitude of the delivered reward dropped linearly as *ET* increased (maximum reward at goal time, GT = 7 s). In early stages of training, smaller rewards were delivered for trials with *ET* < 7 s. However, the smallest *ET* value that triggered reward delivery was progressively raised during learning (see Methods). **c)** Schematic description of an error trial. Early *ET*s triggered an extra-running penalty and an audio noise. *Inset*: the duration of the penalty period was 10 s for the shortest *ET*s and fell linearly to 1 s for *ET*s approaching 7 s. **d)** Schematic description of an omission trial (no beam crossing between 1.5 and 15 s). **(b-d)** Note that *ET*s started to be detected 1.5 s after the motor start.

During the first training sessions, animals started most trials in the front of the treadmill, mostly ran in the reward area and interrupted the infrared beam before the GT (Video 1, Figure 2a, top, c, left). Progressively, across training sessions, animals waited longer and after ∼15 sessions, they reliably entered the reward area just after the GT (Figure 2b). Interestingly, for a large majority of animals, this ability of precisely waiting 7 s before entering the reward area was associated with the performance of a stereotyped motor sequence on the treadmill (Video 2, Figure 2a, bottom, c, right). First, animals began each trial in the reward area. Then, when the treadmill was turned on, they remained largely still while being pushed away from the reward area until they reached the rear wall. Finally, after reaching the rear wall, they ran across the treadmill, without pause, and crossed the infrared beam. The percentage of trials for which animals used this motor routine increased during learning (Figure 2d). Even though a strong preference for the reward area was observed for both correct and error trials, the probability to start a trial in the frontal portion of the treadmill was higher for correct trials compared to error trials (Figure 2e), a tendency that developed progressively during training (Figure S1). In addition, if an animal started a trial in the frontal portion of the treadmill, the probability of reaching the back of the treadmill was higher in correct trials than in error trials (Figure 2f), confirming that correct trials were associated with the animals following the wait-and-run routine and effectively reaching the back of the treadmill before running forward toward the reward area. However, a significant fraction of the animals (14/54) did not develop such a strategy (Figure 2c right, Figure S2a). Compared to these animals, those following regularly the wait-and-run routine entered the reward area later, displayed reduced variability and an increased percentage of correct trials (Figure S2b-d). While we cannot exclude that animals from the middle and front groups also used a more subtle stereotyped motor routine not captured by tracking the average body positions along the treadmill length, the above results suggest that following a front-back-front trajectory through the “wait-and-run” routine is the most reliable strategy to accurately enter the reward area just after 7 s.

**Figure 2:**
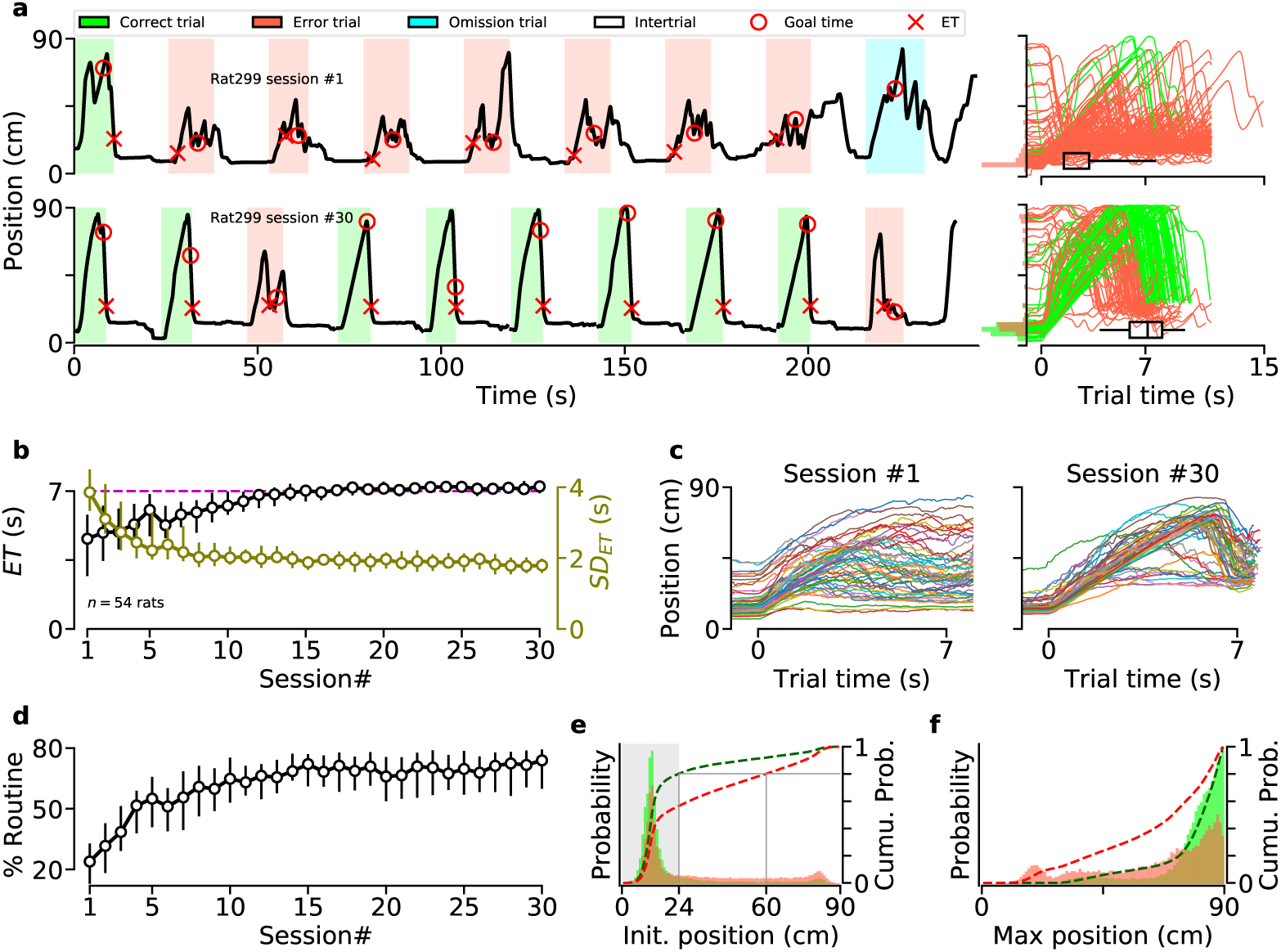
Most animals developed a unique stereotyped motor sequence. **a)** *Left*: illustration of an animal’s trajectory on the treadmill during 9 consecutive trials of the 1st (*top*) and 30th (*bottom*) training sessions. On the y-axis, 0 and 90 indicate the treadmill’s front (reward port) and rear wall, respectively. *Right*: trajectories for all trials during the 1st (*top*) and 30th (*bottom*) sessions (same animal as left panels). Distributions of initial positions for correct (green) and error (red) trials are shown on the y-axis. Black horizontal boxplots depict entrance time range (center line, median; box, 25th and 75th percentiles; whiskers, 5th and 95th percentiles). **b)** Median entrance time (*ET*) in the reward area for the first 30 daily training sessions. Circles indicate group median and error bars, the median range (25th and 75th percentiles) across animals for *ET* and on the right y-axis,*SD* of *ET* (*SD*_*ET*_) values. The dashed magenta line shows the goal time (7 s). **c)** Median trajectory of all the trials for the 1st (*left*) and 30th (*right*) training sessions. Each line represents a single animal (*n* = 54). **d)** Session-by-session percentage of trials during which animals performed the stereotyped front-back-front trajectory (see Methods). Circles indicate group median and error bars, the median range across animals (25th and 75th percentiles). **e)** Probability distribution function (PDF) of the position of the animals at the beginning of each correct (green) and error (red) trial, from sessions #20 to #30. Dashed lines represent cumulative distribution functions (right y-axis). The gray area indicates that in trained animals, 80% of correct trials began with the animal located near the front of the treadmill. **f)** PDF of the maximum position along the treadmill reached by animals before crossing the beam (= *ET*). Only trials in which animals were initially located in the front of the treadmill (gray area in panel e) were included.

It could be argued that a combination of the task parameters (length of the treadmill, its speed and direction, possibility to start trials in the reward area, position of the infrared beam) and the length of the rats’ body (from head to tail) favored the development of this stereotyped strategy. Indeed, depending on the initial position of the animal body at trial onset, it can take up to 7 or 8 seconds for the animals to passively reach the back of the treadmill (Figure 2a) after which they can start running toward the reward area without the need to estimate time. Thus, in the following experiments, we examined how accurately animals respected the GT, when distinct task parameters were modified such as to prevent the use of this simple wait-and-run motor sequence. First, we trained a new group of rats in a version of the task in which, for each trial, the speed of the treadmill was selected randomly from a uniform distribution between 5 and 30 cm/s (Figure 3a). We found that, during the course of training, these animals consistently failed to wait as long as the animals trained in the control version of the task (“control” group, Figure 3b). Still, the average trajectories of animals extensively trained in this “variable speed” condition revealed that they followed a front-back-front trajectory (Figure 3c). Accordingly, the probability of performing a correct trial, given different speeds, fell rapidly from 5 to ∼15 cm/s and was lowest for the fastest treadmill speeds (Figure 3d). Indeed, when the treadmill speed was fast, performing the wait-and-run strategy resulted in error trials, as animals reached the back region of the treadmill earlier than when the treadmill speed was slow. We also found that the probability of entering the reward area at the GT±1 s sharply peaked for a treadmill speed (11.5 cm/s) that is suitable to perform the wait-and-run motor sequence (Figure 3d). Finally, when rats extensively trained in the control version of the task underwent a single probe session with variable speeds (Figure 3e), all measures of performance dropped significantly (Figure 3f). Examining the probability of correct trials and accurate *ET*s (7 ± 1 s) given the treadmill speed suggested that, animals kept performing the wait-and-run routine they previously learned in the control condition (compare Figure 3g and Figure 3d).

**Figure 3:**
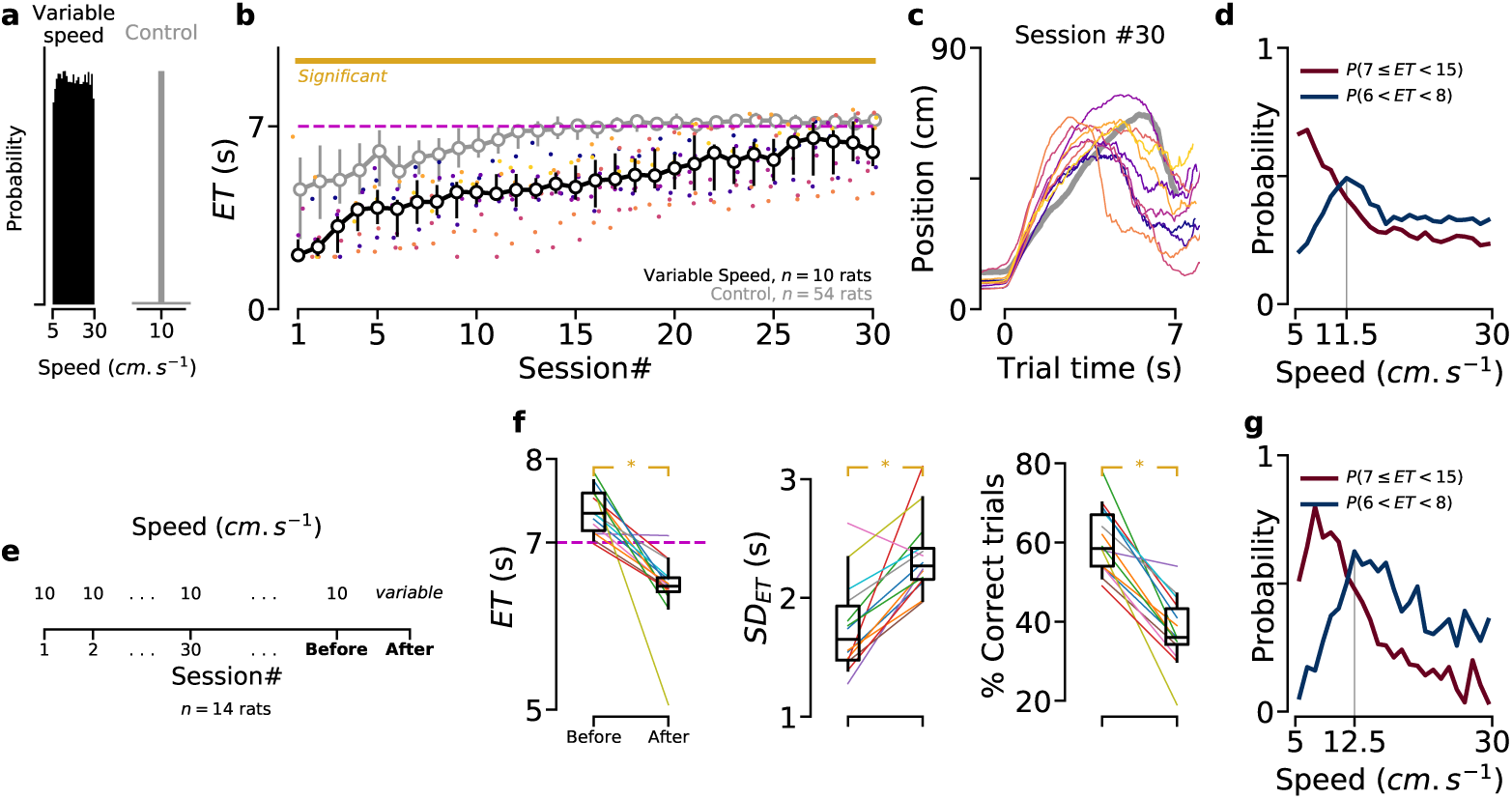
Decreased temporal accuracy when the treadmill speed changes across trials. **a)** For each trial, treadmill speed was either fixed at 10 cm/s (control condition, same data as in Figure 2), or randomly selected from a uniform distribution between 5 and 30 cm/s (variable speed condition). **b)** Median *ET* for animals trained in the variable speed (black), and control (gray) conditions. Colored dots indicate individual performance for “variable speed” animals. Yellow line shows statistically significant differences between groups (permutation test, see Methods). **c)** Median trajectory of “variable speed” animals in session #30 (same colors as in panel b). **d)** Probability of correct (7 ≤ *ET* < 15 *s*) and precise (6 < *ET* < 8 *s*) trial, given the treadmill speed, for “variable speed” animals (session # ≥20). **e)** After extensive training in control condition, animals (*n* = 14) were tested in a probe session with variable speed. **f)** Median *ET*s (*left*), *SD* of *ET*s (*middle*) and percentage of correct trials (*right*) in the sessions immediately before and after the change in speed condition. Each line represents a single animal. Asterisks indicate significant differences (non-parametric paired comparison, see Methods). **g)** Similar to panel d, for the data collected from the probe session.

In the control condition, ∼80% of correct trials started while animals were in the reward area (Figure 2e). If rats relied on an internal clock-based algorithm to accurately time their entrance in the reward area, they should adapt relatively easily to a perturbation of their initial starting position. To test this prediction, we trained a group of rats in a modified version of the task that penalized them when they started the trials in the front region of the treadmill. This was done by activating the infrared beam as soon as the motor was turned on (in the control condition, the infrared beam was inactive during a *timeout* period that lasted 1.5 s after treadmill onset). In this “no-timeout” condition, error trials corresponded to *ET*s occurring between 0 and 7 s after motor onset (Figure 4a). Animals trained in this condition never reached the level of timing accuracy displayed by animals in the control condition (Figure 4b). Still, no-timeout animals followed a front-back-front trajectory (Figure 4c) and correct trials were associated with the animals starting the trials just behind the infrared beam (Figure 4d). The stereotyped reliance on the wait-and-run strategy was also demonstrated by the fact that rats extensively trained in control condition kept performing the exact same trajectory when tested in a single probe session under no-timeout condition, leading to a sharp decrease in performance (Figure 4e-g).

**Figure 4:**
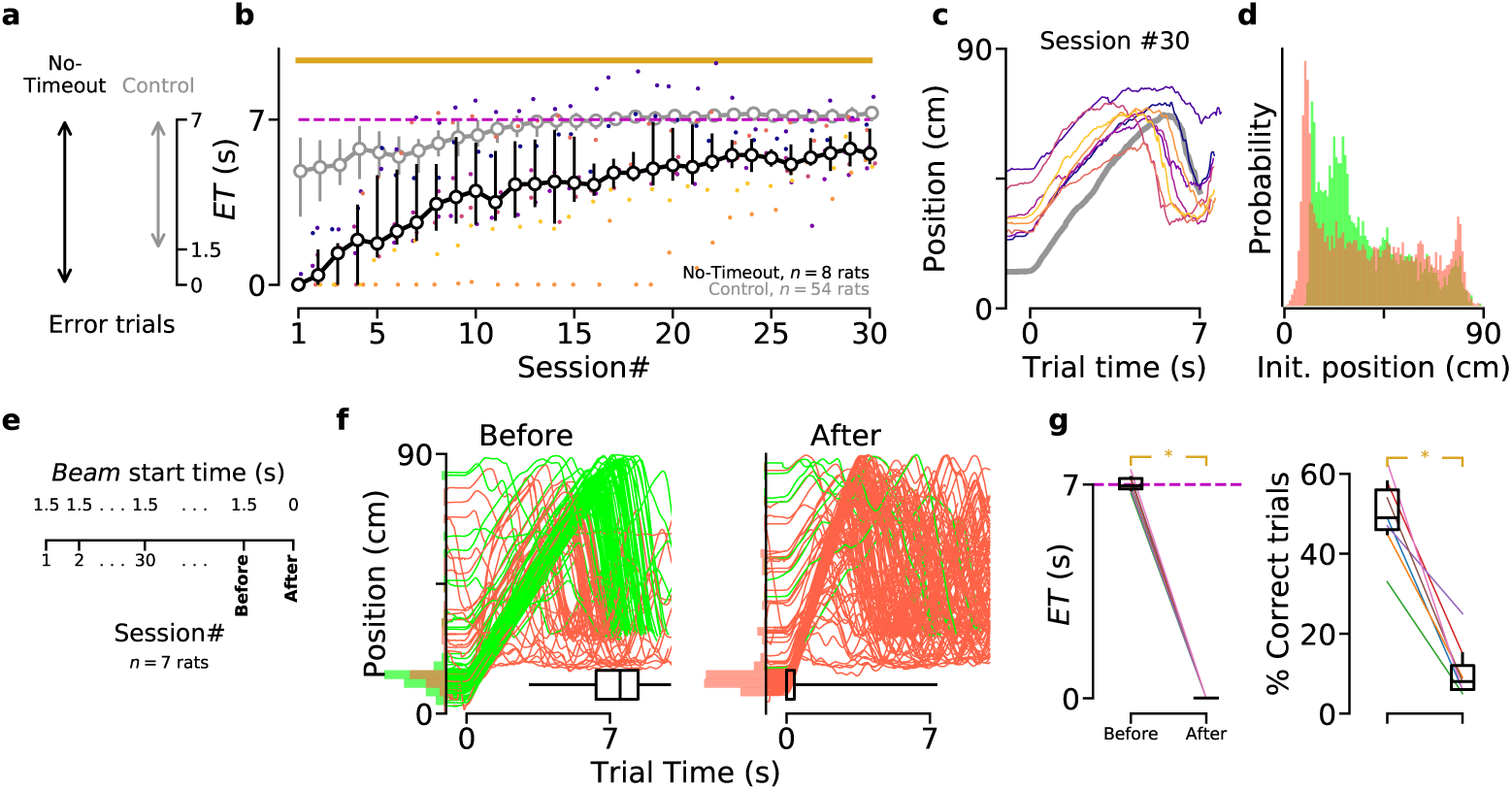
Decreased temporal accuracy when animals are penalized for starting trials in the reward area. **a)** In control condition, animals had a 1.5 s timeout period to leave the reward area after motor onset. In “no-timeout” condition, crossing the infrared beam any time before 7 s is considered as an error. **b)** Median *ET* for animals trained in the no-timeout (black), and control (gray) conditions. Colored dots indicate performance for individual “no-timeout” animals. **c)** Median trajectory of no-timeout animals (same colors as in panel b) in session #30. **d)** PDF of the no-timeout animals’ positions at the beginning of each trial, from sessions #20 to #30. **e)** After extensive training in control condition, animals (*n* = 7) were tested in a no-timeout probe session, in which the beam started at the beginning of the trial, rather than 1.5 s later. **f)** Trajectories of a representative animal in the last “control” session (*left*), and the probe session (*right*). **g)** Median *ET*s (*left*), and percentage of correct trials (*right*) in the sessions immediately before and after the change in beam start time. Each line represents a single animal. Asterisks indicate significant differences (non-parametric paired comparison, see Methods).

We next examined how animals behaved when the goal time (GT) was set to 3.5 s (Figure 5), a condition in which the performance of the wait-and-run strategy would lead to late *ET*s (and smaller rewards) because it can take up to ∼8 s for the animals to passively travel from the front to the rear portion of the treadmill. Animals successfully entered the reward area after 3.5 s and reduced their variability across training sessions (Figure 5a) but as a group, they displayed an increased *ET* variability compared to animals trained in the control condition, with GT set 7 s (Figure 5e). From the averaged trajectories of “short GT” animals measured once their performance plateaued, it appeared that 3 subjects out 7 followed a front-back-front trajectory by running toward the rear portion of the treadmill. The other 4 animals remained still when the treadmill was turned on and tried to run forward before reaching the rear wall (Figure 5b). Interestingly, after training, in 67% of the error trials, the rats started running forward before reaching the middle of the treadmill (Figure 5c, compare with red histogram in Figure 2f). Conversely, after initiating a trial in the reward area, the probability of visiting a deeper portion of the treadmill was much stronger in correct than error trials, reinforcing the idea that accurate timing was accomplished by exploiting the most salient physical features of the environment (Figure 5c). Accordingly, the 3 rats that followed the front-back-front trajectory were less variable than those that passively stayed still before running toward the reward area from the middle of the treadmill (Figure 5e, same color code as in panel b). In addition, among animals trained in the short goal time condition, we found that the magnitude of the backward displacement on the treadmill was negatively correlated with *ET* variability (*r* = −0.49, *p* = 2.7 × 10^−3^, Pearson’s correlation). In the short GT condition, animals became proficient more rapidly than in the control condition (compare Figure 5a with Figure 2c). Could the increased *ET* variance when the GT is 3.5 s be explained by the fact that the task is easier in this condition and that animals do not need to be very precise? To test this possibility, we increased the penalty for early *ET*s and decreased reward size for late *ET*s. In this “sharp reward” condition, the performance of the animals trained in the short GT was even more variable (Figure 5d-e). This result confirms that under short GT condition animals can not accurately time their entrance in the reward area. Finally, another group of animals was trained with GT set to 3.5 s and treadmill speed at 20 cm/s (i.e., twice as fast, such as following the front-back-front trajectory through the wait-and-run motor sequence would lead to *ET*s close to the GT, Figure 5f). These animals displayed reduced *ET* variability compared to animals trained at 10 cm/s, and after treadmill onset they stayed immobile until reaching the end of the treadmill, similar to animals trained in the control condition (Figure 5g,h).

**Figure 5:**
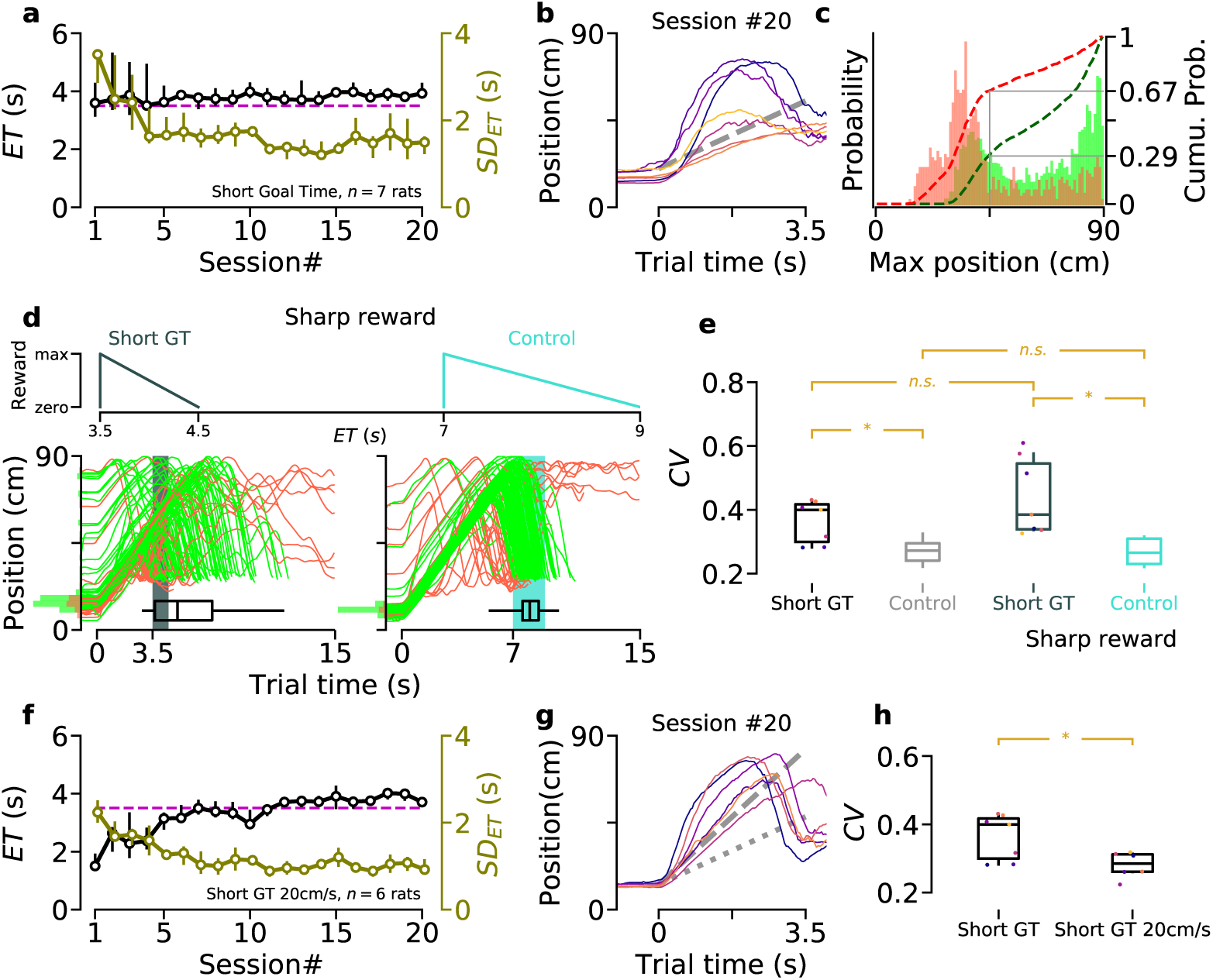
Decreased temporal accuracy when the goal time is shortened. **a)** Median entrance time (*ET*) during training. The dashed magenta line shows the goal time (*GT* = 3.5 s). The right y-axis shows standard deviation (*SD*) of *ET*. **b)** Median trajectory of “short GT” animals after training. Colored lines indicate performance of individual animals. Dashed line’s slope shows the treadmill speed (10 cm/s). **c)** PDF of the maximum position reached by short GT animals before *ET* for correct (green) and incorrect (red) trials. Dashed lines represent cumulative distribution functions (right y-axis). Data collected from session # ≥ 15. **d)** Sharp reward condition applied to short GT and control experiments. *Top*: reward profiles in the sharp condition. *Bottom*: trajectories of 2 illustrative sessions after training in sharp condition (*left*, short GT; *right*, control). Highlighted areas indicate the reward window. **e)** Coefficient of variation (*CV*) for short GT and control experiments with normal (first two boxes), and sharp (last two boxes) reward profiles. Data collected and averaged once performance plateaued (after session #15 for short GT, between session #20 to #30 for control, and last 5 sessions for the sharp condition experiments). Short GT vs. Control: *p* < 0.0001 (permutation test, see Methods); Sharp short GT vs. Sharp control: *p* < 0.0001 (permutation test); Short GT vs. Sharp short GT: Non significant (non-parametric paired comparison); Control vs. Sharp control: *p* = 0.79 (permutation test). **f)** Similar to panel a, for another group of animals that were trained to wait for 3.5 s while the speed of the treadmill was 20 cm/s. **g)** Similar to panel b, for animals trained in short goal time 20 cm/s condition (panel f). Dashed line’s slope shows the treadmill speed (20 cm/s). Dotted line’s slope indicates control treadmill speed (10 cm/s). **h)** *CV* for short GT and short GT 20 cm/s conditions (same colors as in panel b,g). Data collected and averaged once performance plateaued (after session #15). Short GT vs. Short GT 20 cm/s: Asterisk indicates significant difference (10,000 resamples with replacement, see Methods).

The above results suggest that, in a task requiring animal to produce a motor response according to a fixed temporal constraint, the possibility to perform a stereotypical motor sequence adapted to salient features of the environment (here, taking advantage of the full treadmill length and its physical boundaries) critically determines temporal accuracy. To further investigate this idea, we trained a group of animals in a version of the task in which the treadmill was never turned on (trial onset was signalled by turning the ambient light on). In this condition, animals displayed a strong impairment in respecting the GT, compared to animals trained in the control condition (Figure 6a,b). On average, animals reached the reward area later and later across sessions but displayed a constant high variability in *ET* (Figure 6c). We also noticed that correct trials preferentially occurred when animals crossed the treadmill from the rear wall to the reward area (Video 3, Figure 6a,d,e). Accordingly, after extensive training, a robust correlation was observed on a session by session basis between the percentage of correct trials and displacement of the animal on the treadmill (Figure 6f). Moreover, the probability of a correct trial increased for higher values of displacement (Figure 6g).

**Figure 6:**
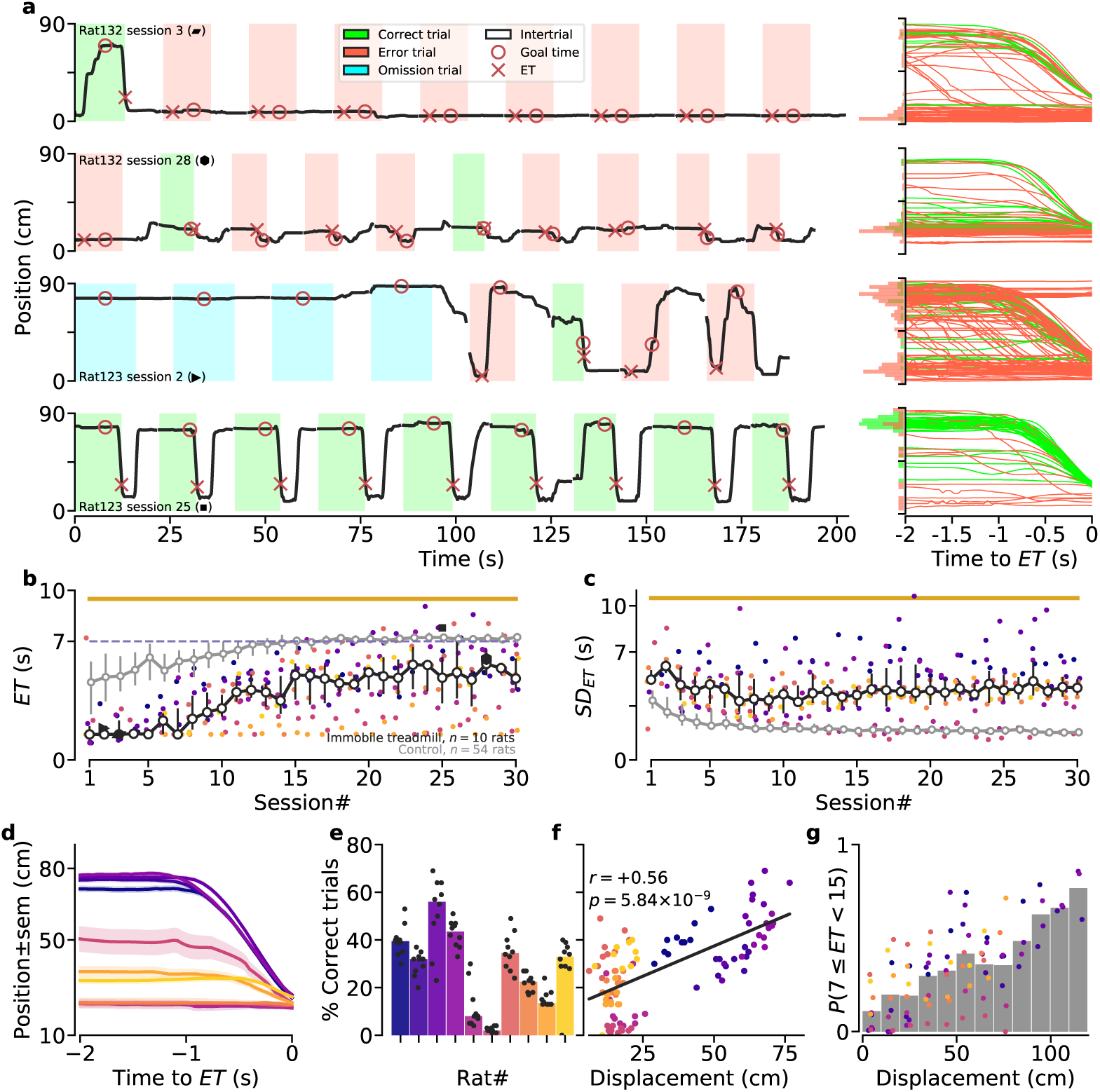
Performance of animals trained while the treadmill remained immobile. **a)** *Left*: illustrations of the positions of two animals on the immobile treadmill for 9 consecutive trials, early (*1st row:* Rat #132-session #3, *3rd row:* Rat #123-session #2) and late (*2nd row:* Rat #132-session #28, *4th row:* Rat #123-session #25) during training. *Right*: trajectories for all the trials of the corresponding sessions on the left, aligned to the *ET*. Distributions of positions 2 s before *ET*, for correct (green) and error (red) trials are shown on the y-axis. Median *ET* across sessions for “immobile treadmill” animals. Filled black markers correspond to the sessions illustrated in panel a. **c)** Similar to panel b, for the standard deviation of entrance times (*SD*_*ET*_). **d)** Median trajectory aligned to *ET* of each “immobile treadmill” animal (only correct trials from sessions #20 to #30 are considered; shaded regions denote standard error). **e)** Median percentage of correct trials for each immobile treadmill animal (same sessions as in panel d). Each dot represents one session. **f)** Repeated measures correlation between the percentage of correct trials and average displacement during a session. Each dot represents one session. **g)** PDF of a correct trial, given the displacement of an animal. Each dot represents the average probability for an individual animal, during a single session. **(e-g)** Analyses based on the same sessions as in panel d. Individual animal color code is preserved in panels b-g.

Lastly, animals trained in the immobile treadmill condition during several weeks were challenged in the control condition (i.e., by simply setting the treadmill speed at 10 cm/s). These animals improved their behavior at the same pace and with the same wait-and-run routine as naïve animals (Figure 7a-c). Thus, animals that previously learned to wait in one version of the task did not learn faster than naïve animals when challenged in a second version of the task with distinct movement requirement but identical GT, demonstrating again that task proficiency relied primarily on the acquisition of a motor sequence rather than an internal knowledge of time.

**Figure 7:**
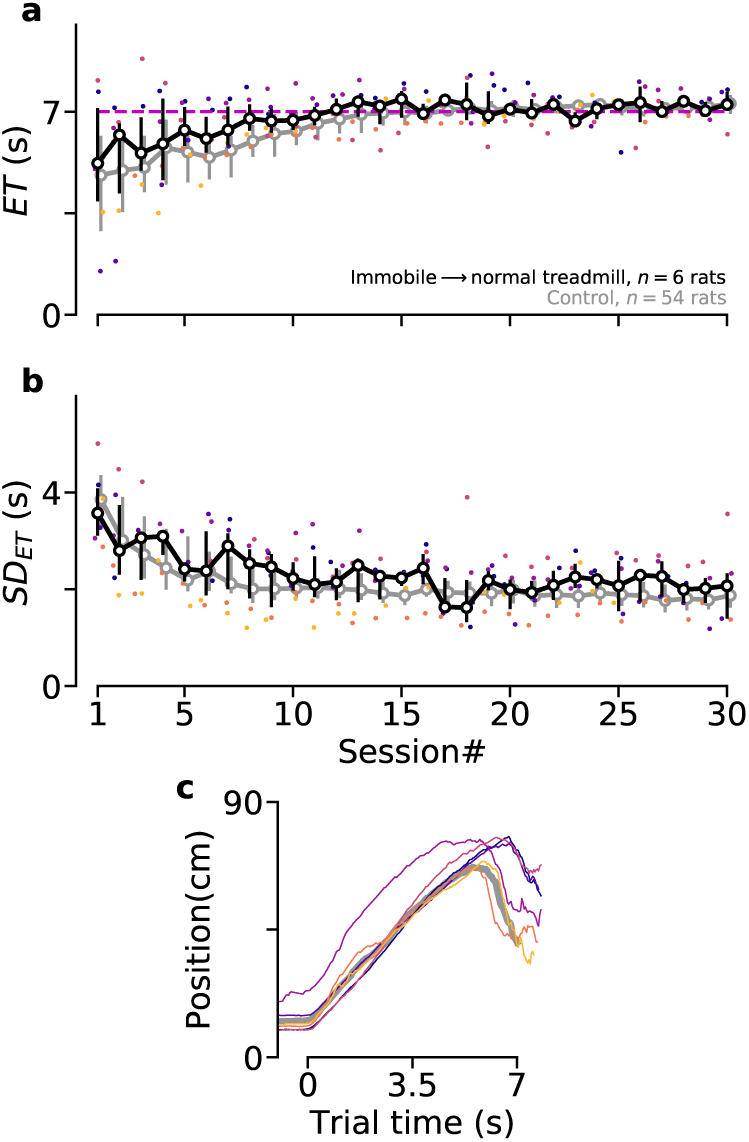
Lack of temporal knowledge transfer across task protocols. After extensive training on the immobile treadmill, animals were trained under normal conditions (GT= 7 s, treadmill speed= 10 cm/s). **a)** Median *ET* across sessions in control condition. **b)** Similar to panel a, for the standard deviation of entrance times (*SD*_*ET*_). **c)** Median trajectory of the individual animals after relearning the task in the control condition. **a-c)** Individual animal color code is preserved in all panels.

## Discussion

In this study, we used a treadmill-based behavioral assay in which rats, once a trial started, where required to wait for 7 s before approaching a reward location. Objectively, animals may accurately time their approaches using either one of the following two mechanisms. First, they may rely on a purely *internal* mechanism (self-sustained neuronal dynamics read by their motor system). In that case, behavioral accuracy should be largely independent of variations in *external* factors (e.g., the speed of the treadmill, the animals position on the treadmill at trial onset, …). In addition, animals would probably stay close to the reward area for most of the duration of the trial (Figure S3d-g). Alternatively, animals may discover by trial-and-error a motor routine adapted to the apparatus and task parameters, whose completion would take them into the reward area at the right time. In that case, timing accuracy should be associated with the stereotyped performance of the routine and should also depend on the structure of the environment that must favor the routine expression (Figure S3a-c, Figure S4).

We report that to accurately wait 7 seconds before approaching the reward port, most rats develop the following “wait-and-run” motor routine. First, rats waited for the beginning of each trial close to the reward area. Then, after trial onset, they remained relatively still while the treadmill carried them toward the back of the treadmill. Finally, when they reached the treadmill’s rear wall, they ran straight to the reward port. Importantly, even for proficient animals, the probability of performing a correct trial was almost null when they started a trial in the back region of the treadmill. In addition, when animals started a trial in the reward area, performing a correct trial was almost exclusively associated with the animals reaching the back portion of the treadmill. Crucially, following extensive training in the control condition, when we modified the task parameters to penalize the stereotyped performance of the wait-and-run routine, the behavioral proficiency and accuracy of the animals dropped dramatically. These results support the hypothesis that, in our task, performing the motor routine is necessary for accurate timing.

It could be argued that the parameters of the task (waiting time, treadmill’s length, speed and direction) provided the ideal condition for the discovery of this simple wait-and-run routine. In other words, in conditions that do not favor the usage of a simple motor routine, rats may time their reward approaches by relying on an internal representation of time that arises from the ability of recurrent neural networks to generate self-sustained time-varying patterns of neural activity [13]. However, we found that in such conditions, rats did not seem capable of accurately timing their entrance in the reward area. First, we trained a group of animals while the treadmill speed randomly changed across trials. Compared to animals trained in the control condition, those trained with a variable speed were less accurate. Additionally, these animals also attempted to use the front-back-front trajectory as shown by an increased probability of correct trials when the treadmill speed allowed it. Second, we trained a different group of rats in a version of the task that penalized them when they started the trials in the reward area. In this condition, solving the task is not possible using the wait-and-run routine since they would generate early entrance times in the reward area. Rats trained in this condition displayed strong accuracy impairment and they kept trying to develop a modified front-back-front trajectory by starting the trials as close as possible to the infrared beam. In all the above experiments, during trials, the treadmill pushed the animals away from the reward area which favors the usage of the wait-and-run routine. To avoid this possible bias, in another experiment, we trained a group of rats on an immobile treadmill. Rats’ performance was poor in this condition, with some animals failing to show any signs of learning. Furthermore, animals that did eventually learn, performed a modified motor sequence during which they ran to the back, performed some movements in the back of the treadmill (that we could not quantify with our video tracking system) and then rushed back to the front. Altogether, we conclude from this set of experiments that rats were unable to use a purely internal representation of time, but always attempted to develop a motor routine in the confined space of the treadmill, routine whose execution duration amounted to the time they needed to wait. This conclusion was also supported by the fact that animals were less accurate in timing their entrance in the reward area when the goal time was set to 3.5 s, compared to the control goal time (7 s). Indeed, in this short goal time condition, the wait-and-run strategy is not optimal, as animals would enter the reward area too late. Thus, the decreased timing accuracy might be explained by the difficulty to “self-estimate” when to start running forward without the help of a salient sensory cue (such as touching the back wall). In support of this idea, in 67% of the error trials, the rats started running forward before reaching even the middle of the treadmill. In addition, a few animals trained in the short goal time condition developed a new stereotyped motor sequence (running to the rear, and back to the front). Interestingly, their entrance times were less variable than animals that remained immobile after trial onset and tried to estimate when to run forward in the middle portion of the treadmill.

It could also be argued that rats never understood our task as a time estimation challenge and this is why they tried to solve it using a motor strategy. While we agree that animals solved the task as a motor learning problem, we believe this type of consideration is irrelevant because our study aimed at testing an objective non-trivial question based on the popular view that animals can use an internal representation of time to adapt their behavior to fixed temporal constraints. This logic was confirmed using a simple simulation of the task in the reinforcement-learning theory framework (Figure S3) and thus, our main experimental result was not predictable before doing the experiments. In addition, to the best of our knowledge, models of timing that emphasize the importance internal processes do not require animals to be explicitly aware of time [7, 13]. Finally, there will always be a limit to the inferences that an experimenter can make regarding the mental state of a behaving animal or its internal model of a task. Even in choice tasks explicitly designed to require rats to estimate the duration of sensory cues, it is unclear if they do so (whatever this might mean for them) and video quantification supported the hypothesis that the animals’ performance in this type of task was also dependent on an overt timing strategy (see below).

A more practical limitation of our work is whether its conclusion is relevant beyond the specifics of our experimental protocols (a supra-second long motor timing task in which the rewarding action is a full-body movement in space). Interestingly, in a study in which rats had to perform two lever presses interleaved by 700 ms, each animal slowly developed an idiosyncratic motor sequence (e.g., 1# first press on the lever with the left paw; 2# touching the wall above the lever with the right paw; 3# second press on the lever with the left paw), lasting precisely 700 ms [23]. The large inter-individual variability reported in this study may arise from multiple possibilities of simple action sequences that can be squeezed in such a short time interval, taking advantage of the proximity of the front wall and lever. Nevertheless, this study provides an additional example in which virtually all animals developed a motor strategy, even if, compared to our task, the time interval was much shorter (< 1 s) and the terminal operant response was distinct (a single lever press). More remarkably, in one of the rare studies that continuously recorded and quantified the full body dynamics of rats performing a sensory duration categorization choice task, it was reported that animals developed highly stereotyped motor sequences during presentation of the sensory cues and that perceptual report of the animals could be predicted by these motor sequences [22]. This result is reminiscent of an earlier study showing that the prediction of rats’ temporal judgement (a 6 s long versus a 12 s long luminous signal) was always better if based on the collateral behavior performed by the animal at the end of the signal than if based on time [25]). Thus, in such temporal discrimination tasks, a stereotyped sequences of movements (collateral behavior) might serve as an external clock and the choice of the animals might be primarily determined by what the animal is doing when a sensory cue disappears rather than by an internal estimation of the duration of that cue. Altogether, these studies support the idea that animals resort to motor strategies to adapt to temporal constraints in a wide range of timing tasks. The novelty of our work is, first, to demonstrate that even in conditions that discourage the use of such motor strategies rats do not seem able to rely on a purely internal timing mechanism and, second, that a critical determinant of temporal accuracy is the possibility to develop motor routines that can be guided by interactions with salient features of the environment.

It has been previously proposed that timing could be mediated through motor routines whose precise execution is internally controlled [18–21]. Thus, it could be argued that accurate timing in our task was ultimately driven by internal neuronal dynamics. We don’t dispute the fact that neuronal activity is required for proficient performance in our task. Actually, we have previously reported that striatal inactivation decreased timing accuracy in a slightly modified version of this task [24]. In addition, there is no reason why the moment-to-moment movement dynamics of the animals on the treadmill could not be decoded from spiking activities recorded across cortical and subcortical regions. However, this type of result can not be used as a definitive evidence in favor of a neuronal representation of time read by the animals as we, humans, watch a clock [26–28]. Indeed, here we report that timing accuracy was reduced when the task parameters prevented the animals from taking advantage of the physical structure of the treadmill to learn the motor routine. Thus, in our task, something more than an internal process (be it a dedicated clock or the self-sustained population dynamics emerging from recurrently connected circuits) was required for accurate timing: the reciprocal and repetitive interactions between the nervous system and the body (sensors and actuators) on the one hand, and the surrounding environment on the other hand. Our results are compatible with the idea that timing emerges from the dynamics of neural circuits [11, 12], as long as these dynamics are not entirely internally generated but also reflect feedback from the environment. For instance, we would assume that the timing deficits induced by striatal inactivation [24] might be explained by the role of this brain region in accumulating sensory information before taking a decision [29, 30].

That timing could be primarily embodied and situated might seem counterintuitive with our innerly rooted feeling of time. Nevertheless, it is interesting to note that to precisely measure time, we have created devices that indicate time by moving objects in space and extensively use metaphors containing movement and space references when speaking of time (“holidays are approaching”, “time flies”) [31, 32]. Moreover, humans display poor temporal judgment accuracy when prevented to count covertly or overtly [33] and several studies have reported that movements improve the perception of rhythmical intervals [34–36] It has been recently proposed that the explicit perception of time in humans may be constructed implicitly through the association between the duration of an interval and its sensorimotor content [37]. The fact that motor timing may be fundamentally related to movement in space for both animals and humans could explain why brain regions involved in movement control and spatial representation, such as the motor cortex, basal ganglia, cerebellum and hippocampus, have consistently been associated with time representation [38–46]. Still, why animals and humans seem to favor embodied and interactive timing strategies over purely internal mechanisms is not clear. Insights regarding this question might be obtained by considering adaptive behavior in an evolutionary perspective [47] and time in the context of ecologically valid timing tasks [48].

## Methods

### Subjects

Subjects were male Long-Evans rats. They were 12 weeks old at the beginning of the experiments, housed in groups of 4 rats in temperature-controlled ventilated racks and kept under 12 h–12 h light/dark cycle. All the experiments were performed during the light cycle. Food was available *ad libitum* in their homecage. Rats had restricted access to water while their body weights were regularly measured. A total of 111 rats were used in this study (the number of animals in each experimental condition is systematically shown in its respective figure). No animal was excluded from the analysis. All experimental procedures were conducted in accordance with standard ethical guidelines (European Communities Directive 86/60 - EEC) and were approved by the relevant national ethics committee (Ministère de l’enseignement supérieur et de la recherche, France, Authorizations #00172.01 and #16195).

### Apparatus

Four identical treadmills were used for the experiments. Treadmills were 90 cm long and 14 cm wide, surrounded by plexiglass walls such that the animals were completely confined on top of the treadmill. Each treadmill was placed inside a sound-attenuating box. Treadmill belt covered the entire floor surface and was driven by a brushless digital motor (BGB 44 SI, Dunkermotoren). A reward delivery port was installed on the front (relative to the turning direction of the belt) wall of the treadmill and in case of a full reward, released a ∼80 *µ*L drop of 10% sucrose water solution. An infrared beam was installed 10 cm from the reward port. The first interruption of the beam was registered as entrance time in the reward area (*ET*). A loudspeaker placed outside the treadmill was used to play an auditory noise (1.5 kHz, 65 db) to signal incorrect behavior (see below). Two strips of LED lights were installed on the ceiling along the treadmill to provide visible and infrared lighting during trials and intertrials, respectively (see below). The animals’ position was tracked via a ceiling-mounted camera (Basler scout, 25 fps). A custom-made algorithm detected the animal’s body and recorded its centroid as animal’s position. The entire setup was fully automated by a custom-made program (LabVIEW, National Instruments). Experimenter was never present in the behavioral laboratory during the experiments.

### Habituation

Animals were handled 30 m per day for 3 days, then habituated to the treadmill for 3 to 5 daily sessions of 30 min, while the treadmill’s motor remained turned off and a drop of reward was delivered every minute. Habituation sessions resulted in systematic consumption of the reward upon delivery.

### Behavioral Task (Normal Condition)

#### Treadmill Waiting Task

Training started after handling and habituation. Each animal was trained once a day, 5 times a week (no training on weekends). Each of the daily sessions lasted for 55 min and contained ∼130 trials. Trials were separated by intertrial periods lasting 15 s. During intertrials, the treadmill remained in the dark and infrared ceiling-mounted LEDs were turned on to enable video tracking of the animals. Position was not recorded during the last second of the intertrials to avoid buffer overflow of our tracking routine and allow for writing to the disk. The beginning of each trial was cued by turning on the ambient light, 1 s before motor onset. Since animals developed a preference to stay in the front (i.e., close to the reward port), the infrared beam was turned on 1.5 s after trial start. This *timeout* period was sufficient to let the animals be carried out of the reward area by the treadmill, provided they did not move forward. After the first 1.5 s, the first interruption of the beam was considered as *ET*. The outcome of the trial depended solely on the value of the *ET*, compared to the goal time (GT= 7 s, except for the short goal time condition). In a correct trial (*ET* ≥ *GT*, see Figure 1), infrared beam crossing stopped the motor, turned off the ambient light, and triggered the delivery of reward. In an error trial (*ET* < *GT*, see Figure 1), there was an extended running penalty for a duration determined by the rule displayed in Figure 1c, inset. During the penalty, the motor kept running, the ambient light stayed on and an audio noise indicated an error trial. In omission trials wherein animals didn’t cross the beam in 15 s since the motor start, trial stopped and reward was not delivered.

#### Reward Profile

The magnitude of the reward was a function of the *ET* and animal’s performance in previous sessions (only in early training, see Figure 1b, inset). Reward was maximal at *ET* = *GT* and dropped linearly to a minimum (= 38% of the maximum) for *ET*s approaching 15 s (i.e., the maximum trial duration). Moreover, in the beginning of the training, partial reward was also delivered for error trials with *ET* > *ET*_0_, where *ET*_0_ denotes the minimum threshold for getting a reward. The magnitude of this additional reward increased linearly from zero for *ET* = *ET*_0_, to its maximum volume for *ET* = *GT*. In the first session of training, *ET*_0_ = 1.5 s and for every following session, it was updated to the maximum value of median *ET*s of the past sessions. Once *ET*_0_ reached the GT, it was not updated anymore (late training reward profile in Figure 1b, inset).

#### Motor Routine

We quantified the percentage of trials in which animals performed the front-back-front motor routine. Trials were considered *routine* if all the following three conditions were met: the animal started the trial in the front (initial position < 30*cm*); 2) the animal reached the rear portion of the treadmill after trial onset (maximum trial position > 50*cm*); 3) the animal completed the trial (i.e., they crossed the infrared beam). The same criteria were applied to the median trajectories after training (session #30) to classify animals into two groups: those that used the front-back-front trajectory and those that did not (Figure S2).

### Alternative Training and Testing Conditions

#### Variable Speed Condition

In this condition, for each trial, treadmill speed was pseudo-randomly drawn from a uniform distribution between 5 and 30 cm/s. During any given trial, the speed remained constant. We used 5 cm/s as the lowest treadmill speed. Lower speeds generated choppy movements of the conveyor belt. Also, velocities higher than 30 cm/s were not used, to avoid any physical harm to the animals.

#### No-timeout Condition

In the control condition, the infrared beam was not active during the first 1.5 s of the trials. This *timeout* period was sufficient to let the animals be carried out of the reward area by the treadmill, provided they did not move forward. In the “no-timeout” condition, the infrared beam was activated as soon as the trial started. Thus, in this condition, error trials corresponded to *ET*s between 0 and 7 s. Consequently, animals were penalized if they were in the reward area when the trial started (i.e., *ET* = 0 s).

#### Short Goal Time Condition

In this condition, the goal time (GT) was set to 3.5 s, half the value for the control condition. The reward profile in this condition followed the same rules as for the control condition, except that reward was maximal at *ET* = *GT* = 3.5 s. Two different groups of animals were trained in this condition, one with treadmill speed set to the normal value of 10 cm/s, and another with treadmill running twice as fast (20 cm/s, see Figure 5). In the short goal time condition, we also examined if the increased variability in *ET* could be attenuated when the penalty associated with early *ET* was increased and when reward magnitude was decreased for late *ET*. This was implemented by doubling the treadmill speed during the penalty period (from 10 cm/s to 20 cm/s), and the reward was delivered for a narrower window of *ET*s (maximal reward at *ET* = *GT* = 3.5 s, and no reward after *ET* = 4.5 s). For proper comparison, we also examined the behavior of rats trained with *GT* = 7 s when the running penalty was increased and the reward was decreased for late *ET*s (maximal reward at *ET* = *GT* = 7 s, and no reward after *ET* = 9 s, see Figure 5d,e).

#### Immobile Condition

In this condition, the treadmill’s motor was never turned on. The ambient light was turned on during the trials and turned off during the intertrials. Error trials were penalized by an audio noise and extended exposure to the ambient light.

### Statistics

All statistical comparisons were performed using resampling methods (permutation test and bootstrapping). These non-parametric methods alleviate many concerns in traditional statistical hypothesis tests, such as distribution assumptions (e.g., normality assumption under analysis of variance), error inflation due to multiple comparisons, and sensitivity to unbalanced group size.

We used the permutation test to compare the performance of two groups of animals during training on a session-by-session basis, such as in Figure 3b, and Figure 4b. To simplify the description (see [49] for more details), let’s assume, as in Figure 3b, we have **X** = [*X*_1_, *X*_2_, …, *X*_*n*_], where *X*_*i*_ is the set of *ET*s of all the animals in session *i*. Similarly, we have **Y** that contains *ET*s from another experimental condition. Here, the null hypothesis states that the assignment of each data point in *X*_*i*_ and *Y*_*i*_ to either **X** or **Y** is random, hence there is no difference between **X** and **Y**.

In short, the test statistic was defined as the difference between smoothed (using Gaussian kernel with *σ* = 0.05) average of **X** and **Y** for each session *i*: *D*_0_(*i*). We then generated one set of surrogate data by assigning *ET* of each animal in session *i* to either *X*_*i*_ or *Y*_*i*_, randomly. For each set of surrogate data, the test statistic was similarly calculated, i.e., *D*_*m*_(*i*). This process was repeated 10,000 times for all the statistical comparisons in this study, obtaining: *D*_1_(*i*), …, *D*_10000_(*i*).

At this step, two-tailed pointwise p-values could be directly calculated for each *i*, from the *D*_*m*_(*i*) quantiles (see [49]). Moreover, to compensate for the issue of multiple comparisons, we defined global bands of significant differences along the session index dimension [49]). From 10,000 sets of surrogate data, a band of the largest *α*-percentile was constructed, such that less than 5% of *D*_*m*_(*i*)s broke the band at any given session *i*. This band (denoted as the *global band*) represents the threshold for significance, and any break-point by *D*_0_(*i*) at any *i* is a point of significant difference between **X** and **Y**.

A similar permutation test was also used when comparing only two sets of unpaired data points (such as in Figure 5e, comparing control vs. short goal time groups). The same algorithm was employed, having only one value for index *i*. If none of the *D*_*m*_(*i*)s exceeded *D*_0_(*i*), the value *p* < 0.0001 was reported (i.e., less than one chance in 10,000).

For paired comparisons (such as in Figure 3f), we generated the bootstrap distribution of mean differences (*n* = 10000 with replacement). Significance was reported (yellow asterisks) if 95% Confidence Interval (CI) of the pairwise differences differed from zero (i.e., zero was not within the CI) [50]. For example, in Figure 3f, right, the 95% CI of pairwise differences is (19, 27)%. Since this interval does not contain zero, it is reported significant, whereas in Figure 5e, the CI of the comparison between normal and sharp short goal time is (−0.17, 0.01) which includes zero, and hence is reported non-significant.

Exceptionally, for the comparison in Figure 5h, even though it is not paired, we used bootstrapping, because we did not have enough data points to perform the permutation test. In this case, the resampled distribution (*n* = 10000 with replacement) for each group was calculated, and it was reported significant, since the distributions did not overlap at 95% CI.

In Figure 6f, we used repeated measures correlation implemented in the Pingouin package [51]. This technique relaxes the assumption of independent data points, since each animal contributes more than one.

### Reinforcement Learning Models

Here, we used the Markov Decision Process (MDP) formalism to analyze how artificial agents learn to perform a simplified version of the treadmill task. According to the MDP formalism, at each time step, the agent occupies a state and selects an action. The probability to transition to a new state depends entirely on the previous action and state, and each transition is associated with a certain reward. The agent tries to maximize future rewards and, in our simulations, we used a simple Q-learning algorithm ([52], see below) to model the way the agent learned an optimal policy (i.e., which action to take for any possible state).

We modeled the treadmill task using a deterministic environment in which the time was discretized and the treadmill was divided in 5 regions of equal length. In this simplified setting, we simulated two types of agents that differed only by the type of the information available to them to select actions and analyzed how their behaviour varied.

The first type of agents did not use an explicit representation of time to perform the task. At each time step *t*, the state *s*_*t*_ (i.e., the information used to select actions) consisted in the agent’s position *p*_*t*_, in the treadmill and in a boolean variable *w*_*t*_, whose value was equal to 1, if the agent had previously reached the rear wall during the trial and 0, otherwise. Given these assumptions, each state can be written as *s*_*t*_ = {*p*_*t*_, *w*_*t*_} and the state space consisted of 5 pair of states (a total of 10 states).

The second type of agents in addition, benefited from the information on the elapsed time since the beginning of the trial. Thus, each state was represented as *s*_*t*_ = {*p*_*t*_, *w*_*t*_, *t*}.

For both types of agents, the task was simulated in an episodic manner and the initial position *p*_0_ at the beginning of each trial was assigned randomly as follows: the probability *P*(*p*_0_) that the initial state corresponds to *p*_0_ was proportional to 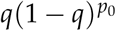 for *p*_0_ = 0, …, 4. We set the parameter *q* = 0.5 such as to account for the tendency of the rats to initiate trials in the reward area.

During the rest of the trial, at each time step *t*, agents occupied a state *s*_*t*_, and could select one of three different actions that determined a transition to a new state *s*_*t*+1_. Action *a*_*t*_ = 0 corresponded to remaining still and, considering that the treadmill was on, moving one position backward on the treadmill. Action *a*_*t*_ = 1 consisted in moving at the same speed of the treadmill (*v*_*T*_), but in the opposite direction. Thus after performing this action, the agents remained at the same position on the treadmill. Finally, performing action *a*_*t*_ = 2, the agents moved at twice the treadmill speed which made him move one position step forward. We also introduced two physical constraints that limited the action space at the extreme sides of the treadmill. In the front of the treadmill, the agents cannot move forward (i.e., when the position was *p* = 0 the action *a* = 2 was forbidden). In the rear of the treadmill the agents could not stay still, as otherwise it would hit the rear wall (i.e., when *p* = 4 the action *a* = 0 was not available).

After entering a new state at time *t* + 1, the agents received a reward 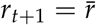. The value 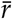 varied depending on the position *p*_*t*+1_ and on the current time *t*. Similarly than in the real task, the agent had to reach the most frontal region of the treadmill (equivalent of the reward area) after 7 time steps (the minimum *ET* in the frontal region to obtain a reward is 8 time steps). We also created an equivalent of the time out period (see above in experimental method section), such as the agent was not penalized to start a trial in the reward area. Still, the agents had to leave the front of the treadmill (i.e., *p* = 0) within 2 time steps. Finally, agents had a maximum amount of time (15 time steps) to perform the task. More specifically, reward rules were as follows. The punishment associated with an early *ET* (2 ≤ *ET* < 8) had a maximum (negative) value of 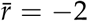 and its absolute value decreased linearly between 2 and 7. Correct trials occurred when agents reached the frontal region of the treadmill between 8 and 15 time step (8 ≤ *ET* ≤ 15), which delivered a reward with a maximum value of 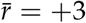, that decreased linearly with *ET*. Omission trials (i.e., those trials in which the agent did not approach the front area within 15 time steps) were associated with the delivery of a small punishment 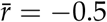. We also modeled the cost of the passage of time while the treadmill was on, by adding a small punishment 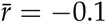 at each time step in all trial types.

Agents learned the value (expressed in terms of future rewards) of selecting a particular action in a specific internal state via the Q-learning algorithm. Specifically, for any state-action pair {*s, a*}, a state-action value function *Q*(*s, a*) can be defined as follows:

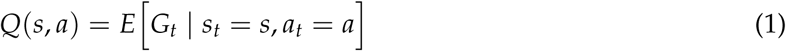

where 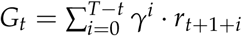 is the discounted sum of expected future rewards, and *γ* is the discount factor (0 ≤ *γ* ≤ 1). Equation 1 implies that each value *Q*(*s, a*) is a measure of the future reward that the agent expects to receive after performing action *a* when its current state is *s*.

Following the Q-learning algorithm, after each time step *t*, the *Q*(*s*_*t*_, *a*_*t*_) will change according to:

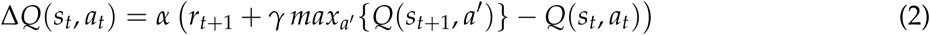

where the parameter *α* represents the learning rate.

These state-action values are then used to determine the policy *π*: a mapping from states to actions (i.e., the way agents acted in any possible state). In our model, the policy was stochastic and depended on the Q-values via a *softmax* distribution:

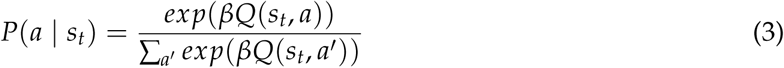

where the parameter *β* governs the exploitation/exploration trade-off (when *β* → 0, the policy becomes more and more random).

Updates in Equation 2 can be proved to converge to the optimal Q-value for each pair {*s, a*} [52]. Optimal value means the value (in terms of rewards) that action *a* assumes in state *s*, when the policy of agent across all the sequence of states and actions is such to maximize future rewards. Therefore selecting actions with a probability that increases with the Q-values allows learning of the optimal behavior.

We used the formalism described above to simulate *n* = 15 agents of the first type and *n* = 15 of the second type. Each agent differed in the exploitation/exploration parameter (see below) and performed the task for 30 sessions of 100 trials each. The exploitation/exploration parameter started with an initial value *β*_0_, and was increased after each session of training by an amount Δ*β* (i.e., the policy became more and more greedy), up to a maximum of *β*_*max*_ = 10. Different agents were represented by different values of *β*_0_ and Δ*β*. The agents of our simulations corresponded to all the possible combinations of *β*_0_ = {0, 2, 2.5, 3, 4} and Δ*β* = {0.3, 0.35, 0.4}. In all the simulation, we set the parameters *α* = 0.1, and *γ* = 0.99.

### Data Analysis

Data from each session was stored in separate text files, containing position information, entrance times, treadmill speeds, and all the task parameters. Position information was scaled to the treadmill length, and smoothed (Gaussian kernel, *σ* = 0.3 s). The entire data processing pipeline was implemented in python, using open-source libraries and custom-made scripts. We used a series of Jupyter Notebooks to process, quantify, and visualize every aspect of behavior, to develop and run the reinforcement learning algorithms, and to generate all the figures in this manuscript. All the Jupyter Notebooks, as well as the raw data necessary for full replication of the figures and videos are publicly available via the Open Science Foundation (https://osf.io/7s2r8/?view_only=7db3818dcf5e49e88d708b2597a21956).

## Author contributions

Mostafa Safaie: Conceptualization, Software, Visualization, Investigation, Formal analysis, Writing– original draft; Maria-Teresa Jurado-Parras: Conceptualization, Investigation, Writing–review & editing; Stefania Sarno: Software, Methodology, Formal analysis, Conceptualization, Visualization; Jordane Louis: Investigation; Corane Karoutchi: Investigation; Ludovic F. Petit: Methodology, Resources; Matthieu O. Pasquet: Methodology, Resources, Software; Christophe Eloy: Conceptualization, Methodology; David Robbe: Funding acquisition, Project administration, Supervision, Visualization, Writing–original draft, Conceptualization, Software.

## Acknowledgements

We thank Drs. I. Bureau and J. Epsztein for critical reading of the manuscript. This work was supported by European Research Council (ERC-2013-CoG – 615699 – NeuroKinematics; D.R.) and a Centuri postdoctoral fellowship (S.S.).

## Supplementary Material

**Video 1.** Video clip showing several consecutive trials from an animal performing its first training session in control condition. Information about trial number, time since light on, GT, *ET*, and ongoing task status are given on the upper left corner.

**Video 2.** Same as Video 1 for a well-trained animal performing the task in control condition.

**Video 3.** Same as Video 2 for an animal performing the task in the immobile treadmill condition.

The 3 videos can be seen and downloaded via the Open Science Foundation (https://osf.io/7s2r8/?view_only=7db3818dcf5e49e88d708b2597a21956)

**Supplementary Figure S1:**
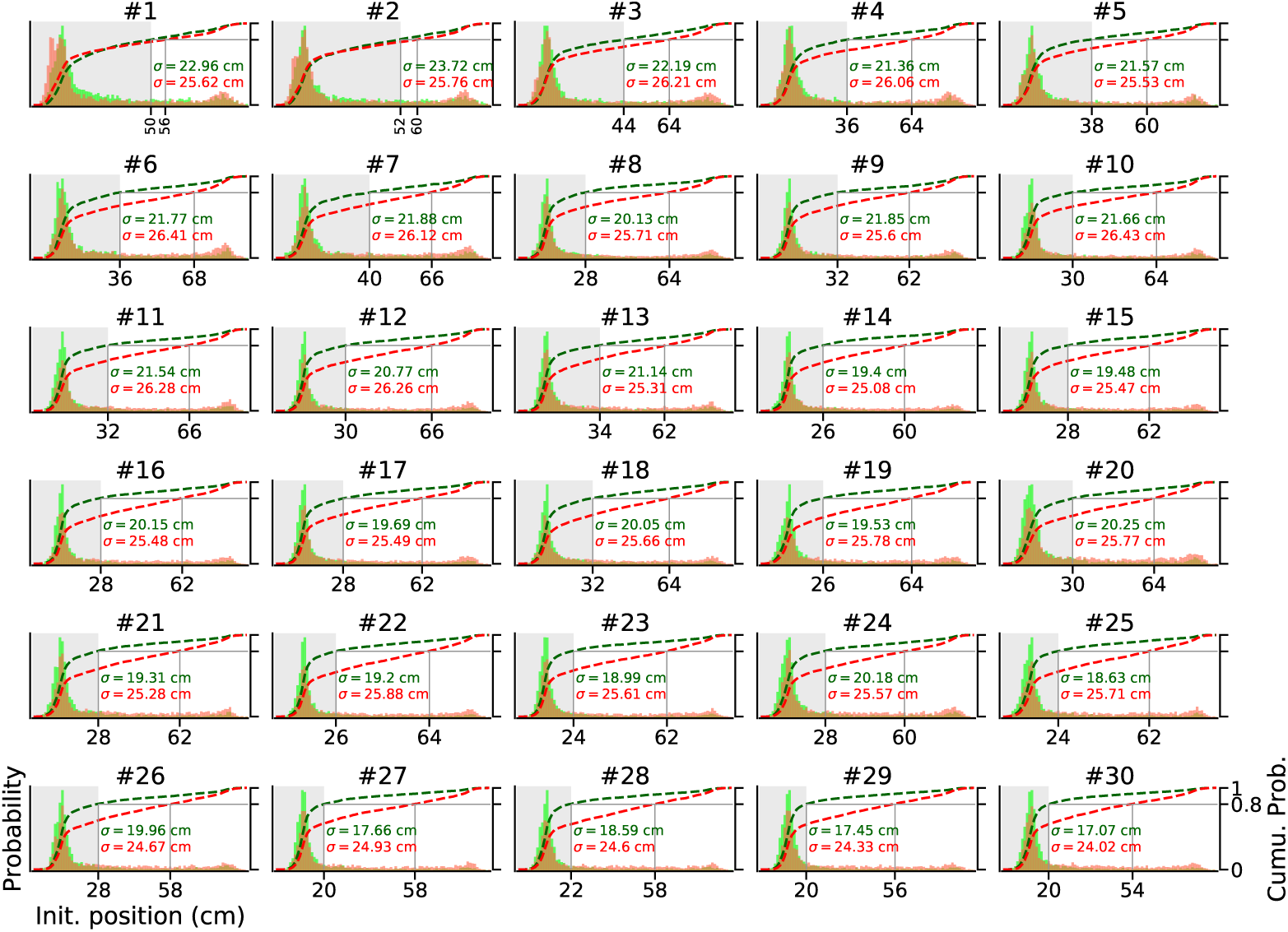
Initial position distributions for correct and error trials diverged progressively during training. Similar to Figure 2e, each panel shows PDF of the initial position of the animals for correct (green) and incorrect (red) trials, but plotted separately for each training session (#1 to #30). Dashed lines represent cumulative distribution functions (right y-axis). For each PDF, *σ* values denote the standard deviation. Each PDF included pooled data from all the animals trained in the control condition (*n* = 54).

**Supplementary Figure S2:**
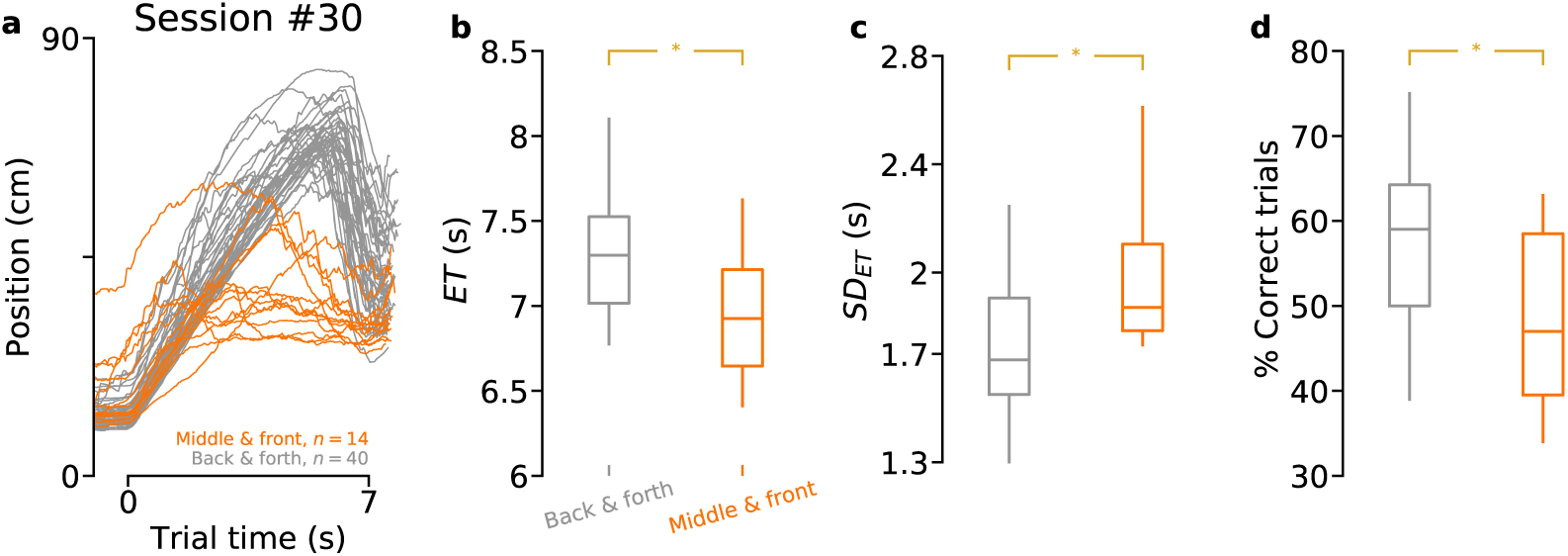
Task proficiency according to the type of trajectory performed by animals. **a)** Same as Figure 2, panel c, right, but the animals were divided in two groups according to whether they performed the front-back-front trajectory (gray) or not (other, orange). **b)** Entrance times (*ET*s). *p* = 0.0066 (permutation test). **c)** *SD* of *ET. p* = 0.03 (permutation test). **d)** Percentage of correct trials. *p* = 0.01 (permutation test). For panels b, c, d, same color code as in panel a. Data from sessions # ≥ 20 were averaged for each animal.

**Supplementary Figure S3:**
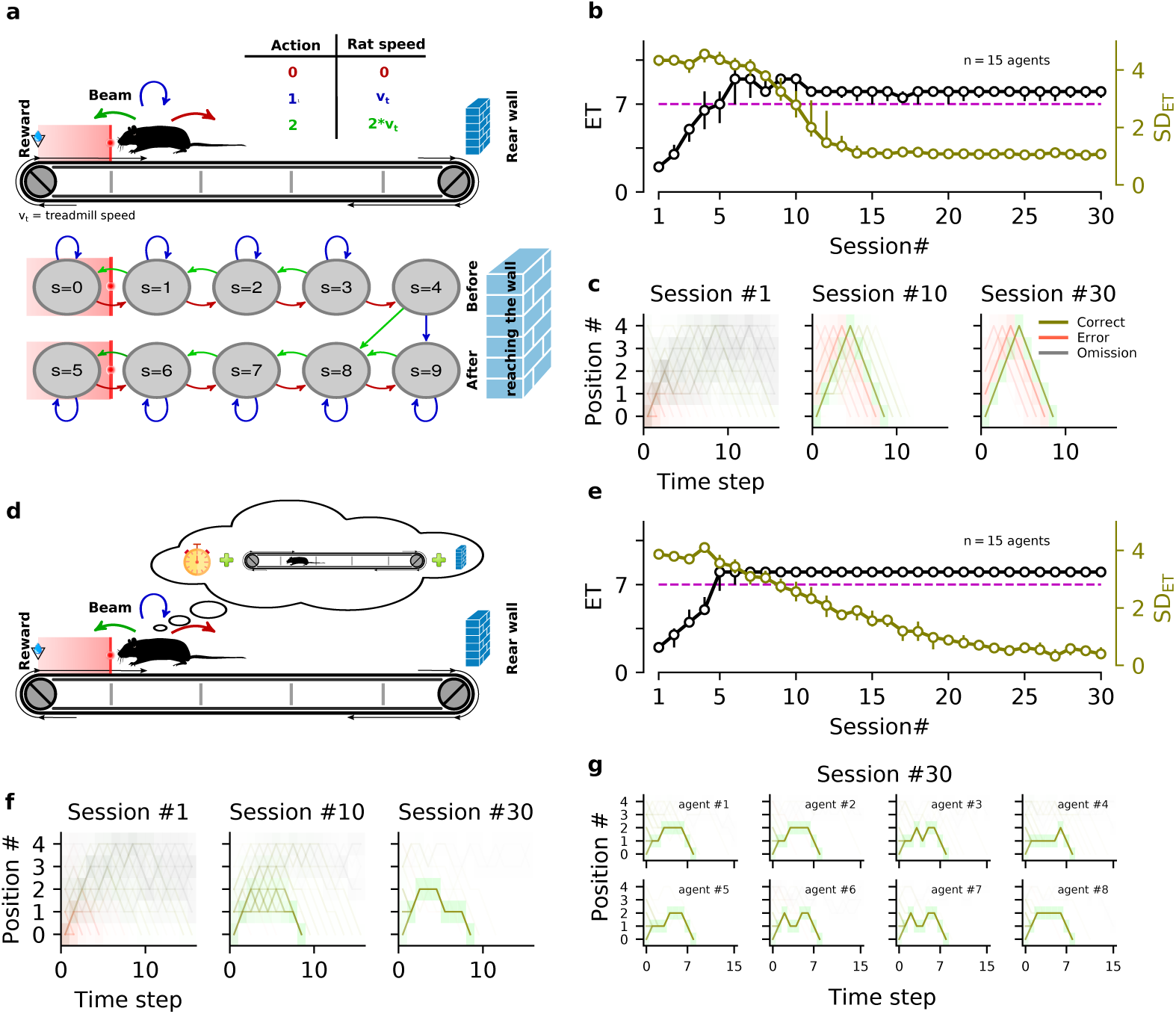
Performance comparison between artificial agents with or without time knowledge. **a)** Schematic representation of the state-action space and transitions used to model the treadmill task for agents that did not use time information. Space and time were discretized. The same physical location corresponded to 2 different internal states before and after reaching the rear wall. At each time step, agents could choose one of the three following actions: staying still (red arrow, action *a* = 0); moving at the treadmill speed *v*_*T*_ (blue arrow, *a* = 1); or moving at twice *v*_*T*_ (green arrow, *a* = 2, see Methods for details). **b)** Average learning profile for several agents with different learning parameters (see Methods). Median entrance time (*ET*) for the first 30 sessions. Error bars show *ET* range (25th and 75th percentiles). The dashed magenta line shows the goal time (7 time steps). The right y-axis shows standard deviation (*SD*) of *ET*. **c)** Trajectories of three sessions at different stages of learning. Each session contained 100 trials. For each session, the space-time occupancy matrix was normalized to its maximum value, for correct (green), error (red), and omission (gray) trials. Across training sessions, the artificial agents (simulated rats) waited longer and longer and reduced their entrance time variability by performing the front-back-front trajectory. The same strategy was developed by agents endowed with different learning parameters and exploration/exploitation rates (Figure S4). After learning, the agents performed the front-back-front routine independently of their initial positions. Thus, they arrived in the reward area too early when their initial position was near the middle of the treadmill, in a striking similarity with the behavior of well-trained rats. **d)** Same as in panel a, but agents have now access to the time elapsed since trial onset. **e-f)** Same as panels b-c, respectively, but for agents following the model sketched in panel d. Agents with temporal knowledge also learned progressively to enter in the reward area at the right time and reduced their variability. However, after learning, they did not perform the front-back-front trajectory. Rather, they mainly stayed behind the infrared beam and were capable of respecting the GT independently of their initial position. **g)** Trajectories after training for 8 different agents that accessed time information. The different agents developed idiosyncratic position trajectories (even if they remained close to the reward area during most of the trial duration), due to stochastic variations in their learning dynamics.

**Supplementary Figure S4:**
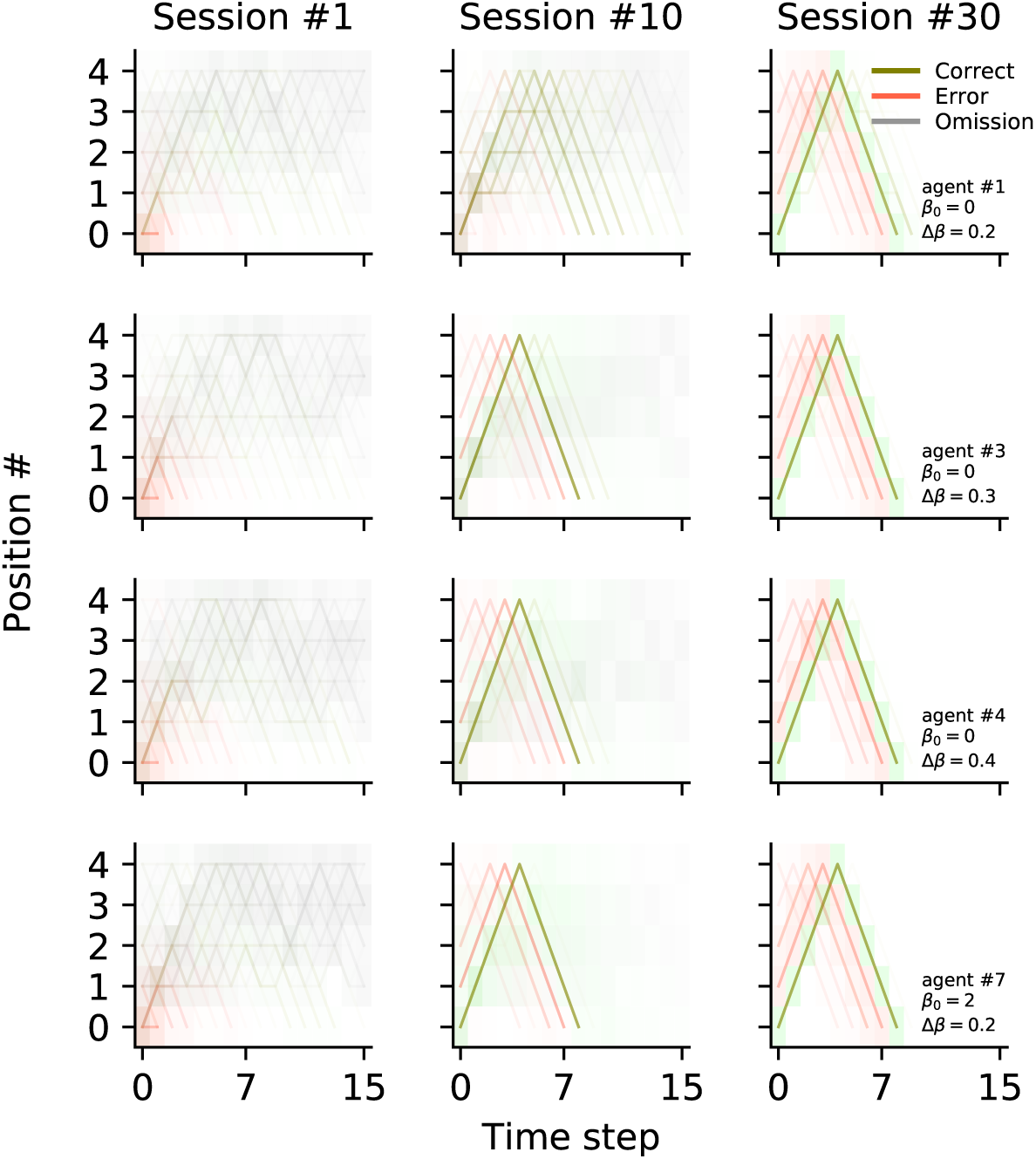
Final trajectories performed by agents are identical regardless of exploitation/exploration parameters. Similar to Figure S3c but for four different agents (differences among agents are determined by the values of the exploitation/exploration parameters *β*_0_ and Δ*β*; see Methods). Even if agents displayed different trajectories during learning (sessions #1 and #10), all of them performed the same trajectory at session #30.

